# Questioning unexpected CRISPR off-target mutations in vivo

**DOI:** 10.1101/157925

**Authors:** Sang-Tae Kim, Jeongbin Park, Daesik Kim, Kyoungmi Kim, Sangsu Bae, Matthias Schlesner, Jin-Soo Kim

## To the Editor

Recently, Schaefer et al.^1^ reported that whole genome sequencing (WGS) of two Cas9-treated, gene-corrected mice and a wild-type control mouse unveiled 1,397 single-nucleotide variations (SNVs) and 117 small insertions and deletions (indels) present commonly in the two Cas9-treated mice “but absent in the uncorrected control” and from a database of mouse SNVs and indels. There was essentially no sequence homology between the on-target site and these SNVs and indel sites, most of which lacked a protospacer-adjacent motif (PAM) sequence, suggesting that these variations were both small guide RNA (sgRNA)-independent and Cas9-independent, respectively. Nevertheless, the authors made a bold claim that these variations were caused by CRISPR-Cas9 without validating these unexpected off-target effects even at a single SNV or indel site by performing an independent experiment in vitro or in vivo. Another major concern in Schaefer et al. is the absence of analysis of variants that are present in the control mice but absent in the two gene-edited mice.

Target specificities of CRISPR systems have been extensively studied in animals and cell lines. For example, we showed that certain CRISPR-Cas9 nucleases did not induce detectable off-target mutations at sites with just two or three-nucleotide mismatches in human cells, first using T7 endonuclease I assays^2^ and then using targeted deep sequencing^3^. We and others also performed WGS to show that Cas9 rarely induced off-target indels in a clonal population of cells^4-7^ or a gene-edited animal^8^. Note that Schaefer et al. did not find any off-target mutations in the two Cas9-treated mice at the top 50 most likely off-target sites with 3- to 4- nucleotide mismatches, in line with these previous reports. Given the remarkable specificity of CRISPR-Cas9, it is difficult to believe that Cas9 can cleave sites that differ by > 10 nucleotides from the on-target sequence, as suggested in Schaefer et al.

The authors did not articulate whether the unexpected off-target effects were limited to the particular target site or their method or FVB/NJ zygotes used in their experiments. In silico off-target predicting algorithms and cell-based, genome-wide off-target profiling methods such as GUIDE-seq^9^ and HTGTS^10^ cannot identify off-target sites with no sequence homology. However, Digenome-seq, an in vitro method of capturing in vitro cleavage sites using Cas9-digested, cell-free genomic DNA via WGS, does not rely on sequence homology^4^, because DNA double-strand break ends remain intact in vitro and are protected from digestion by endogenous exonucleases in vivo. Note that we did not find any unusual, non-homologous off-target sites in human genomic DNA using Digenome-seq^4^, showing that such sites cannot be cleaved by Cas9 in vitro.

It is also suspicious that they found a 12-fold excess of SNVs over indels. *S. pyogenes* Cas9 rarely produces SNVs even at on-target sites. After analyzing our targeted deep sequencing data, we found that Cas9 could induce SNVs, when a single base pair (bp) was deleted at a cleavage site and, at the same time, a single bp was inserted, a very rare event that occurred at frequencies of < 1% at on-target sites. Off-target SNVs can be induced much less frequently. In contrast to Cas9-induced mutations, naturally-occurring genetic variations between siblings or strains are strongly biased towards SNVs rather than indels, further casting doubt on “unexpected mutations”.

Accordingly, we hypothesized that the two gene-corrected, possibly sibling, mice were genetically distant from the control “co-housed” mouse, although Schaefer *et* al. indicated that all three mice might belong to the same inbred FVB/NJ strain, and that the SNVs and indels were not caused by CRISPR-Cas9 but inherited from their parents. To test this hypothesis, we used Strelka^11^ and Mutect^12^ that had been employed by Schaefer et al. to find SNVs which are different between two samples in any pairwise-combination of F03, F05, and FVB. We found the lowest number of sample-specific variants when comparing the two samples F03 and F05 (Fig. 1a), while in comparisons including FVB the number of sample-specific variants was considerably higher. Strikingly, this was not only the case for variants present in F03/F05 (and not in FVB), but also for variants present in FVB (and not in F03/F05). To preclude a calling bias caused by the different sequencing depth of F03/F05 and FVB, we used the multi-sample variant caller Platypus^13^ to analyze shared and sample-specific variants (Fig. 1b and 1c). This analysis confirms that FVB bears a similar number of variants which are not present in F03/F05 as vice versa. Furthermore, most variants not present in FVB are shared between F03 and F05, and are homozygous in both of the samples. It is virtually impossible that any mutagenic process causes such a pattern. Finally we focused on the heterozygous variants in the three mice. Mice from inbred strains are generally not expected to contain substantial numbers of heterozygous variants, whereas mutagenic processes will in the vast majority cause heterozygous mutations. While there are indeed heterozygous variants in F03 and F05 which are sample-specific or shared between those samples, the number of heterozygous variants is higher in FVB. The largest fraction of heterozygous variants is shared between all three mice, proving that these variants are inherited and not individually acquired in the investigated animals. Altogether, these results demonstrate that there is considerable genetic heterogeneity between the three mice, with F03 and F05 being genetically closer to each other than to FVB.

**Fig. 1.**
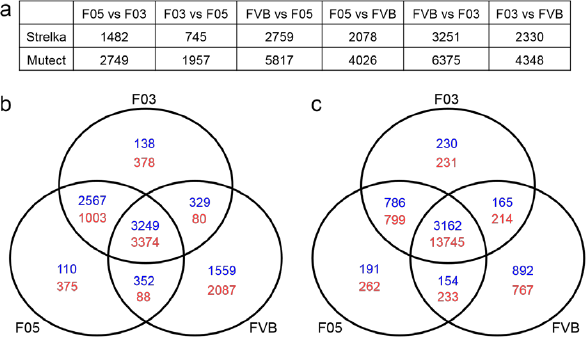
(a) Sample-specific SNVs in all pairwise combinations F03, F05, and FVB (A vs B means variants present in B and not in A). All three aligned sequencing data representing WT (SRR5450998), F03 (SRR5450997), and F05 (SRR5450996) were retrieved and analyzed using Strelka and Mutect. All SNVs were filtered using dbSNP 143 and mouse genome project v5. (b,c) Venn diagrams showing homozygous (blue) and heterozygous (red) SNVs (b) and indels (c) found in the wild-type control mouse (FVB) and the two gene-edited mice (F03 and F05). Variants were identified with Platypus (v. 1.0.8) with multi-sample calling mode with default parameters; variants with coverage lower than 15 in any of the samples, and multi-allelic variants are not considered.

In summary, the CRISPR-Cas9-treated mice do not exhibit more variants that are absent in the uncorrected control than vice versa. Furthermore, the observed variants are extremely unlikely a consequence of a mutagenic process in the investigated animal, but are much better explained by differences in the genetic background between the two gene-edited mice and the control mouse. We conclude that the data presented by Schaefer et al does not provide any evidence for off-target mutagenesis due to CRISPR-Cas9 treatment and see no reason to be doubtful about the previous findings that Cas9 does not cause SNVs and indels at non-homologous, PAM-free sites.

## Acknowledgements

J.-S.K. is supported by a grant from the Institute for Basic Science (IBS-R021-D1).

## Supplementary Note

The SRA files of three mice (FVB mouse: SRR5450998, F03 mouse: SRR5450997, F05 mouse: SRR5450996) were retrieved from NCBI and Bam files were extracted from the SRA files. Mutect 1.1.5 and Strelka 1.0.15 with default options were used for differential (somatic) variant calling using all possible pair-wise combination from the three mice. Platypus 0.8.1 with default options was used for multi-sample variant calling. For all cases only high quality variants were considered for the statistics (i.e. "PASS” in filter column), and variants with coverage threshold below 15 were not considered. All SNVs and Indels found in dbSNP 146 or Mouse genomes project v5 were filtered out using in-house VCF file processing scripts.

## References

1. Schaefer, K.A. et al. Unexpected mutations after CRISPR-Cas9 editing in vivo. Nat. Methods 14, 547 (2017).

2. Cho, S.W., Kim, S., Kim, J.M., & Kim, J.-S. Targeted genome engineering in human cells with the Cas9 RNA-guided endonuclease. Nat. Biotechnol. 31, 230–232 (2013).

3. Cho, S.W. et al. Analysis of off-target effects of CRISPR/Cas-derived RNA-guided endonucleases and nickases. Genome Res. 24, 132–141 (2014).

4. Kim, D. et al. Digenome-seq: genome-wide profiling of CRISPR-Cas9 off-target effects in human cells. Nat. Methods. 12, 237–243 (2015)

5. Smith, C. et al. Whole-genome sequencing analysis reveals high specificity of CRISPR/Cas9 and TALEN-based genome editing in human iPSCs. Cell Stem Cell 15, 12–13 (2014).

6. Veres, A. et al. Low incidence of off-target mutations in individual CRISPR-Cas9 and TALEN targeted human stem cell clones detected by whole-genome sequencing. Cell Stem Cell 15, 27–30 (2014)

7. Suzuki, K. et al. Targeted gene correction minimally impacts whole-genome mutational load in human-disease-specific induced pluripotent stem cell clones. Cell Stem Cell 15, 31–36 (2014)

8. Iyer, V. et al. Off-target mutations are rare in Cas9-modified mice Nat. Methods 12, 479 (2015)

9. Tsai, S.Q. et al. GUIDE-seq enables genome-wide profiling of off-target cleavage by CRISPR-Cas nucleases. Nat. Biotechnol. 33, 187–197 (2015).

10. Frock, R.L. et al. Genome-wide detection of DNA double-stranded breaks induced by engineered nucleases. Nat. Biotechnol. 33, 179–186 (2015).

11. Saunders, C.T. et al. Strelka: Accurate somatic small-variant calling from sequenced tumor-normal sample pairs. Bioinformatics 28(14): 1811–1817 (2012).

12. Cibulskis, K. et al. Sensitive detection of somatic point mutations in impure and heterogeneous cancer samples. Nat. Biotechnol. 31, 213–219 (2013).

13. Rimmer, A. et al. Integrating mapping-, assembly– and haplotype-based approaches for calling variants in clinical sequencing applications. Nat. Genetics 46, 912–918 (2014)

